# circRAB3IP modulates cell proliferation by reorganizing gene expression and mRNA processing in a paracrine manner

**DOI:** 10.1101/2022.03.25.485787

**Authors:** Natasa Josipovic, Karoline K. Ebbesen, Anne Zirkel, Adi Mackay-Danieli, Christoph Dieterich, Leo Kurian, Thomas B. Hansen, Argyris Papantonis

**Affiliations:** Institute of Pathology, University Medical Center Göttingen, 37075 Göttingen; Department of Molecular Biology and Genetics (MBG), Aarhus University, Aarhus, Denmark; Interdisciplinary Nanoscience Centre (iNANO), Aarhus University, Aarhus, Denmark; Center for Molecular Medicine Cologne, University of Cologne, 50931 Cologne; Bioinformatics and Systems Cardiology, University Hospital Heidelberg, 69120 Heidelberg, Germany; Institute of Neurophysiology, University of Cologne, 50931 Cologne

**Keywords:** circRNA, cell proliferation, isoform switching, exosomes, cell-cell communication, senescence

## Abstract

Circular RNAs are endogenous long-lived and abundant non-coding species. Despite their prevalence, only few circRNAs have been dissected mechanistically to date. Here, we catalogued nascent RNA-enriched circRNAs from primary human cells and functionally assign a role to circRAB3IP in sustaining cellular homeostasis. We combined “omics” and functional experiments to show how circRAB3IP depletion deregulates hundreds of genes, suppresses cell cycle progression, and induces senescence-associated gene expression changes. Conversely, excess circRAB3IP delivered to endothelial cells via extracellular vesicles suffices for accelerating their division. We attribute these effects to the interplay between circRAB3IP and the general splicing factor SF3B1, which impacts transcript usage of cell cycle-related genes. Together, our findings link the maintenance of cell homeostasis to the tightly regulated titers of a single circRNA.

## INTRODUCTION

The question of non-coding RNA authenticity and, in many cases, functionality has generally been settled via advances in the technologies developed to catalogue and study transcriptomes. However, due to their structure and unconventional biogenesis (Jeck & Sharpless, 2014; Xiao *et al*, 2020), circular RNAs (circRNAs) remained long overlooked in transcriptome studies. circRNAs are robustly detected across the evolutionary tree (Jeck & Sharpless, 2014; Wu *et al*, 2020), yet their expression is cell type- (Salzman *et al*, 2013), tissue- (Maass *et al*, 2017; Xia *et al*, 2017) and context-specific (Farooqi *et al*, 2021; Knupp & Miura, 2018; Venø *et al*, 2015; Lee *et al*, 2019b). It then follows that circRNA expression is highly regulated, advocating for their functional relevance *in vivo*.

It was only recently that circRNAs emerged as important players in numerous cellular processes, including development and cell differentiation (Lee *et al*, 2019a; Di Agostino *et al*, 2020), senescence (Du *et al*, 2016), immunity (Zhou *et al*, 2019b), cancer (Panda, 2018), Alzheimer’s disease (Zhao *et al*, 2016) and diabetes (Braicu *et al*, 2019). Mechanistically, most of the functions ascribed to circRNAs to date involve their interaction with microRNAs or RNA binding proteins (RBPs) (Du *et al*, 2016, 2017; Abdelmohsen *et al*,2017; Xia *et al*, 2018; Schneider *et al*, 2016). Thus, circRNAs have been shown to act as “sponges” sequestering specific microRNAs or as “decoys” sequestering RBPs. Via such circRNA-microRNA/-RBP interactions the targets of sequestered microRNAs or RBPs are affected as regards their expression (Hansen *et al*, 2013; Zhao *et al*, 2016; Li *et al*, 2015a), RNA processing (Zhang *et al*, 2013; Li *et al*, 2015c; Chen *et al*,2018) or translation (Ashwal-Fluss *et al*, 2014; Abdelmohsen *et al*, 2017). Despite compelling data on the mode-of-action of a handful of circRNAs, how this large class of endogenous non-coding RNAs contribute to cellular homeostasis remains rather poorly understood.

In this study, we investigated circRNAs specifically enriched in the nascent RNA fraction of primary human cells. We focused on the two most enriched cirRNAs in this fraction to show that circRAB3IP can regulate the proliferation potency of cells and, thus, their path to replicative senescence. We also show that circCAMSAP1, earlier implicated in type-2 diabetes (Haque *et al*, 2020), cancer cell physiology (Zhou *et al*, 2020a; Luo *et al*, 2021; Chen *et al*, 2021), and chronic inflammation (Liu *et al*, 2019), also acts as a modulator of senescence-associated inflammatory responses. Notably, circRAB3IP can be packaged into extracellular vesicles and delivered to cells to exert its function. Thus, we provide evidence of how the depletion or paracrine action of a single circRNA to a normal human cell can profoundly remodel its gene expression program and modulate its phenotype.

## RESULTS

### Identification and characterization of circRNAs enriched in nascent RNA

We initially hypothesized that circRNAs with a role in gene regulation would be enriched in nascent RNA. To identify them, we used primary human endothelial cells (HUVECs; pooled from 3 donors) and factory-seq (Melnik *et al*, 2016; **Fig. 1A**). We analysed HUVEC nascent transcriptomes for circRNA enrichment via DCC (Cheng *et al*, 2016) to identify 244 candidates (**Fig. S1A,B**). Of these, only circCAMSAP1 (hsa_circ_ 0001900) and circRAB3IP (hsa_circ_0000419) showed a robust circRNA/mRNA enrichment (DCC > 0.5; **Fig. 1B**) and were supported by detection and high DCC ratios in total RNA-seq data (**Table S1**). Moreover, both were not present in chromatin-associated RNA isolated during factory-seq (**Fig. S1A** and **Table S1**).

**Figure 1.**
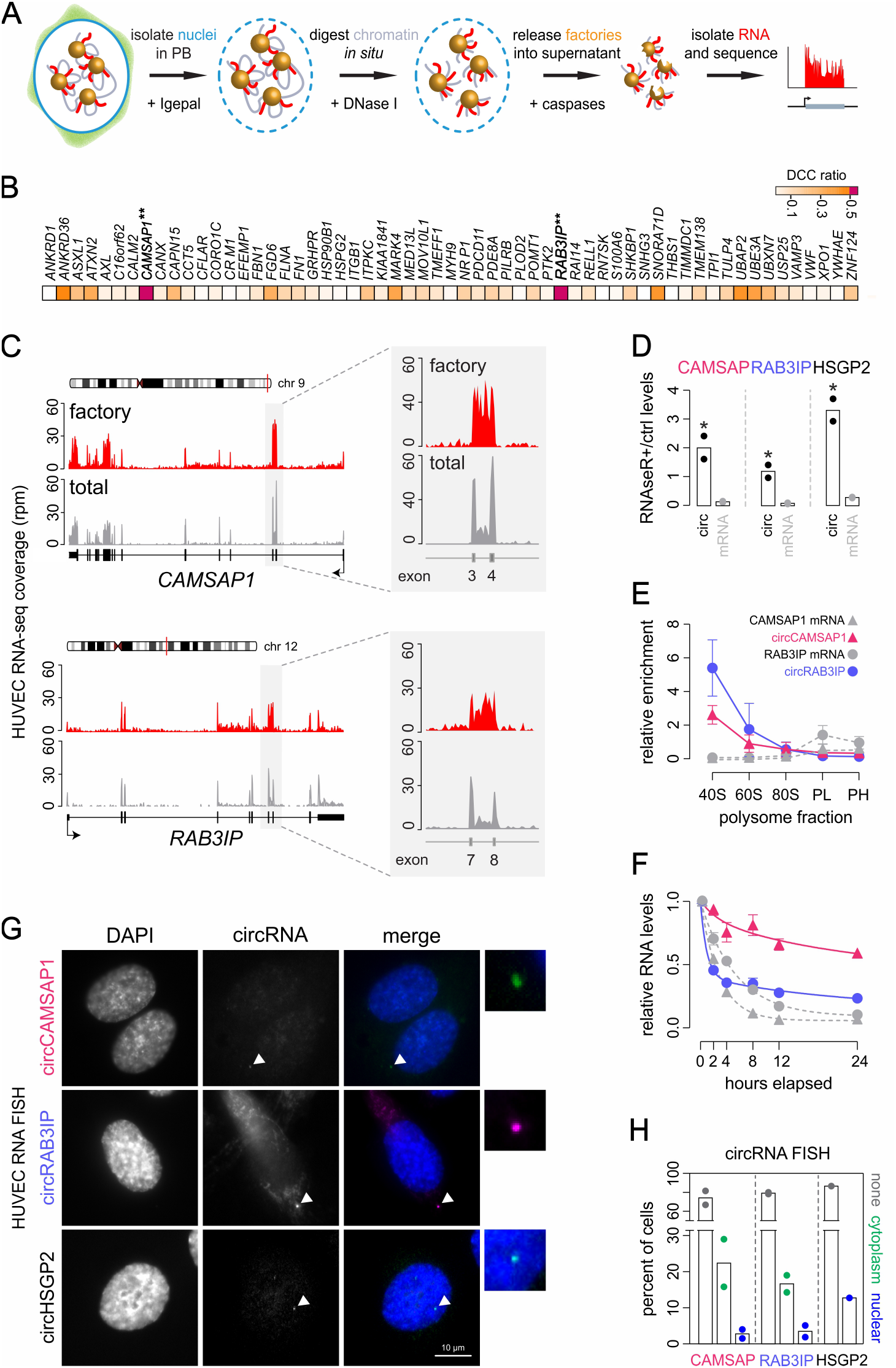
circRNAs enriched in nascent RNA fractions. **A,** Nascent RNA (*red*) from HUVECs is collected following nuclei purification, and DNase I and Group-III caspase treatment to detach transcription factories (*gold*) from the substructure. **B,** Heatmap showing circRNA enrichment over their linear counterparts in factory-seq data represented as a DCC ratio. circRNAs with a ratio >0.5 are highlighted (*starred*). **C,** Factory/total RNA-seq coverage along *CAMSAP1* and *RAB3IP*. Signal enrichment between exons is highlighted. **D,** Bar graph showing mean fold enrichment (from two replicates) of RNase R-treated over untreated samples as determined by RT-qPCR. *: *P*<0.05; unpaired two-tailed Student’s t-test. **E,** Relative enrichment of circRNAs and mRNA counterparts in polysome fractions normalized to whole-cell levels (±SD, from three replicates) determined by RT-qPCR. PL: light polysomes; PH: heavy polysomes. **F,** Decay plots of the RNAs from panel E following transcriptional inhibition for 0-24 h. RNA levels (±SD, from three replicates) were normalized to *YWHAZ* and plotted relative to 0-h levels. Half-lives were: CAMSAP1_mRNA_ = 2.0 h, circCAMSAP1 = 6.2 h, RAB3IP_mRNA_ = 3.7 h, circRAB3IP = 1.0 h. **G,** Representative circCAMSAP1 and circRBA3IP FISH signals (*arrowheads*) in HUVEC counterstained with DAPI. Signal from the non-nascent RNA-enriched circHSGP2 provides a control. Bar: 10 μm. **H,** Bar plots showing percent of cells (from two experiments) without focal circRNA FISH signal (*gray*), with at least one cytoplasmic (*green*) or at least one nuclear focus (*blue*) in images like those in panel G.

circCAMSAP1 and circRAB3IP arise from the sequences of *CAMSAP1* exons 3 and 4, and *RAB3IP* exons 7 and 8, respectively (**Figs 1C** and **S1C**). Characteristic signal enrichment in the intron between the two circRNA-exons (Yang *et al*, 2011) was observed for *CAMSAP1* and *RAB3IP* in factory-seq, but not in total RNA profiles despite greater sequencing depth (**Figs 1C** and **S1D**). The prevalent circRAB3IP isoform was bi-exonic (237 nt-long), whereas the two circCAMSAP1 isoforms containing or not the intron between exons 3 and 4 were comparably abundant (**Fig. S1C**). Primers designed for all subsequent experiments do not distinguish between these two circCAMSAP1 isoforms.

Due to their covalently-closed structure, circRNAs are insensitive to RNase R degradation, display half-lives longer than those of their mRNA counterparts (Jeck & Sharpless, 2014), and are rarely translated (Huang *et al*, 2021). Although both circCAMSAP1 and circRAB3IP had been previously detected in HUVECs (http://www.circbase.org/), we confirmed RNase R resistance (**Fig. 1D**) and lack of polysome association (**Fig. 1E**), validating that both are *bona fide* non-coding circRNAs. As regards their stability *in vivo*, we found an expected long half-life for circCAMSAP1, but not for circRAB3IP (**Fig. 1F**), potentially indicative of tighter temporal regulation. We also tested their dependence on ADAR1, an RNA-editing enzyme affecting circRNA biogenesis (Ivanov *et al*, 2015). We used knockdowns and total RNA-seq to identify ADAR-sensitive circRNAs genome-wide (**Fig. S1D-F**). Following ADAR1-KD, both circCAMSAP1 and circRAB3IP showed decreased DCC ratios indicative of reduced circularization (**Fig. S1G** and **Table S1**).

circCAMSAP1 and circRAB3IP enrichment in nascent RNA could imply association with the active sites of transcription (Caudron-Herger *et al*, 2015). However, circRNAs often localize in the cytoplasm (Salzman *et al*, 2013), as does circCAMSAP1 in cancer cells (Zhou *et al*, 2020a). Therefore, we investigated the subcellular distribution of our circRNAs using fluorescently-labelled 50-mers as probes targeting each backsplicing junction in circRNA FISH (Zirkel & Papantonis, 2018). circCAMSAP1 and circRAB3IP exhibited both nuclear and cytoplasmic localization in HUVECs. In ~5% of cells, circRNA signal accumulated focally at the outer border of the nucleus (**Fig. 1G,H**). In contrast, control experiments targeting circHSPG2, an exonintron circRNA with a high DCC ratio but not enriched in nascent RNA (**Figs 1D**, **S1C** and **Table S1**), showed exclusive nuclear localization (**Fig. 1G,H**). We complemented this analysis with RT-qPCR on nuclear or cytoplasmic RNA to find slight (for circCAMSAP1) to high (for circRAB3IP) nuclear enrichment (**Fig. S1H**). This agrees with factory-seq enrichments and implies quick translocation from the nucleus to the cytosol.

### circRNA depletion leads to senescence-like transcriptional profiles

We next asked whether depletion of either circCAMSAP1 or circRAB3IP from HUVECs would alter their gene expression program. siRNA-mediated circRNA-knockdown (KD; **Fig. S2A**) followed by total RNA sequencing showed large-magnitude transcriptional changes with 3134 and 920 differentially expressed genes (DEGs) upon circCAMSAP1- and circRAB3IP-KD, respectively (**Fig. 2A-C**). Genes upregulated upon circCAMSAP1-KD were mostly linked to immune/inflammatory responses, while those downregulated to the regulation of the cell cycle and ribosome metabolism (**Fig. 2D**). Genes downregulated upon circRAB3IP-KD were also predominantly related to cell cycle regulation, as well as to RNA metabolism (**Fig. 2E**). Unlike circCAMSAP1-KD, circRAB3IP depletion did not trigger proinflammatory gene expression (**Fig. 2D,E**). We also analyzed enrichment of transcription factor motifs known to regulate DEGs in each circRNA-KD. Motifs for E2F factors, involved in cell cycle control (Dimova & Dyson, 2005), dominated downregulated genes analysis in both datasets, while motifs of proinflammatory like NF-κB, STATs and IRFs (Platanitis & Decker, 2018) were overrepresented for genes upregulated by circCAMSAP1-KD (**Fig. S2B**).

**Figure 2.**
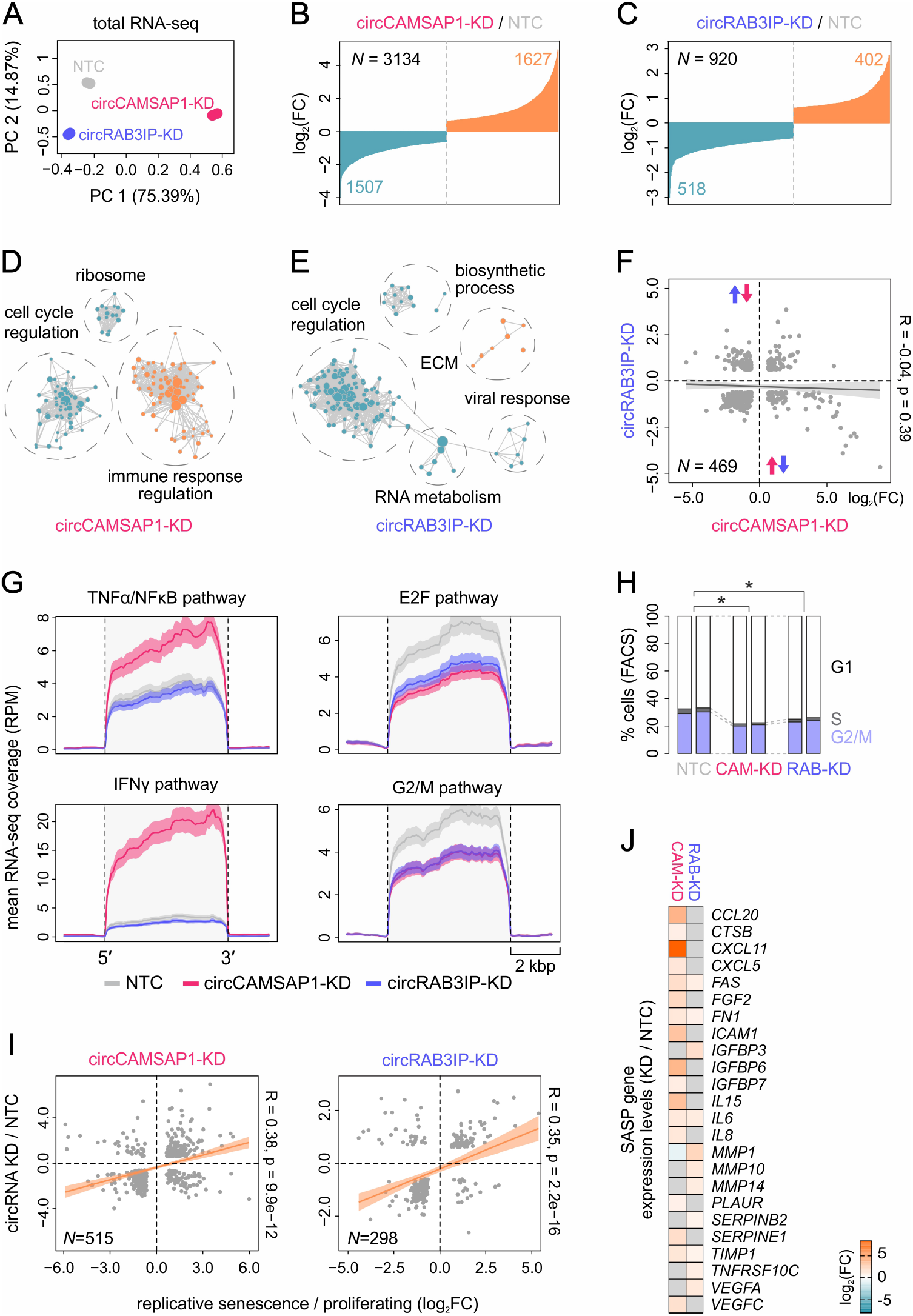
circCAMSAP1- and circRAB3IP-KD trigger major transcriptional changes in HUVECs. **A,** PCA clustering of RNA-seq replicates from circCAMSAP1- (*magenta*), circRAB3IP-KD (*blue*) and non-targeting controls in HUVECs (NTC; *grey*). **B,** Bar plot of fold change (log_2_) for genes up-/downregulated (*orange/blue*) in circCAMSAP1-KD data with a *P*_adj_ ≤ 0.05 and absolute fold change (log_2_) ≥ 0.6 **C,** As in panel B, but for circrRAB3IP-KD data. **D,** Gene set enrichment analysis of DEGs from panel B. Related terms are clustered (*dotted circles*) and labelled by the group-dominant gene ontology term. **E,** As in panel D, but for circRAB3IP-KD DEGs. **F,** Scatter plot correlating fold-change values (log_2_) for the DEGs (*N*) shared by circCAMSAP1- and circRAB3IP-KD. Pearson’s correlation coefficient (R) and its associated p-value are indicated. **G,** Plots of circCAMSAP1- (*pink*) and circRAB3IP-KD (*blue*) mean normalized coverage across all genes relevant to inflammation and cell cycle regulation. NTC data coverage (*grey*) provides a baseline. **H,** Bar plots depicting cell cycle profiles determined via FACS of PI-stained HUVECs in circRNA-KD experiments (two replicates). *: significantly different to NTC; *P*<0.05, Fisher’s exact test. **I,** As in panel F, but correlating circRNA-KD DEGs with those from HUVEC replicative senescence. **J,** Heatmap of SASP genes that are also DEGs in circCAMSAP1- and circRAB3IP-KD data. Color coding reflects foldchange values (log_2_) at a *P*_adj_ ≤ 0.05.

The two circRNA-KD datasets shared >450 DEGs, but the respective changes showed no significant correlation (**Fig. 2F**). Nevertheless, genes downregulated in both experiments were part of E2F-controlled and G2/M cell cycle pathways, and this was reflected in changes of the cell cycle profiles of KD cells showing extended G1 phases (**Fig. 2G,H**). On the other hand, IFNβ/γ and TNFα signaling genes were stimulated by circCAMSAP1-KD only (**Fig. 2G**). Thus, the partially overlapping roles of these two circRNAs unfold via non-convergent regulation of different effector genes.

The cell cycle deceleration that followed both circCAMSAP1- and circRAB3IP-KD, complemented by inflammatory stimulation and ECM remodeling, was reminiscent of the changes accompanying replicative senescence entry by HUVECs (Zirkel *et al*, 2018). Therefore, we compared each KD dataset with DEGs from senescent HUVECs to discover significant correlation (R > 0.35; **Fig. 2I**). In fact, ~30% of circRAB3IP- and 16% of circCAMSAP1-KD DEGs were regulated in the same manner upon senescence entry. Even stronger correlation (R > 0.55) was observed when comparing our KD datasets to a consensus gene expression signature of senescence (Hernandez-Segura *et al*, 2018; **Fig. S2C**), and the few genes showing divergent regulation did not enrich for any particular pathway or GO term. Senescent cells also display cell type-specific secretory phenotypes (i.e., SASP) comprised of molecules acting in a paracrine fashion to promote inflammation and senescence (Coppé *et al*, 2010). SASP genes induced in our circRNA-KD cells (**Fig. 2J**) represent ~1/3 off all relevant entries in the Reactome database (https://reactome.org/). In line with our GSEA analysis (**Fig. 2D,E**), circCAMSAP1-KD induced the largest number of SASP genes. Together, our results indicates that depletion of either circRNA suffices for the extensive remodeling of the HUVEC homeostatic program into senescence-like gene expression programs.

### circRAB3IP modulates the processing of cell cycle-related transcripts

Various studies have suggested that circRNAs regulate cell proliferation via microRNA “sponging” (Zeng *et al*, 2018; Li *et al*, 2018; Zheng *et al*, 2016). However, this requires multiple cognate binding sites for the sequestered microRNA in the circRNA (e.g., as in Hansen *et al*, 2013), which does not appear to be the case for circRAB3IP or circCAMSAP1 (assessed via http://www.targetscan.org). Thus, we set out to explore the association of either circRNA with regulatory proteins. To this end, we performed RNA antisense pull-down coupled to mass spectrometry (RAP-MS, McHugh & Guttman, 2018; **Fig. 3A**) in HEK293 cells. This choice of cell type was dictated by the need for large numbers of cells (4-5 x 10^8^ per replicate) overexpressing circRAB3IP or circCAMSAP1 (**Fig. S3A,B**). Prior to RAP-MS, we performed circRNA-KDs in HEK293 confirming that the majority of the gene expression changes were like those seen in HUVECs (**Fig. S2A,D-G**).

**Figure 3.**
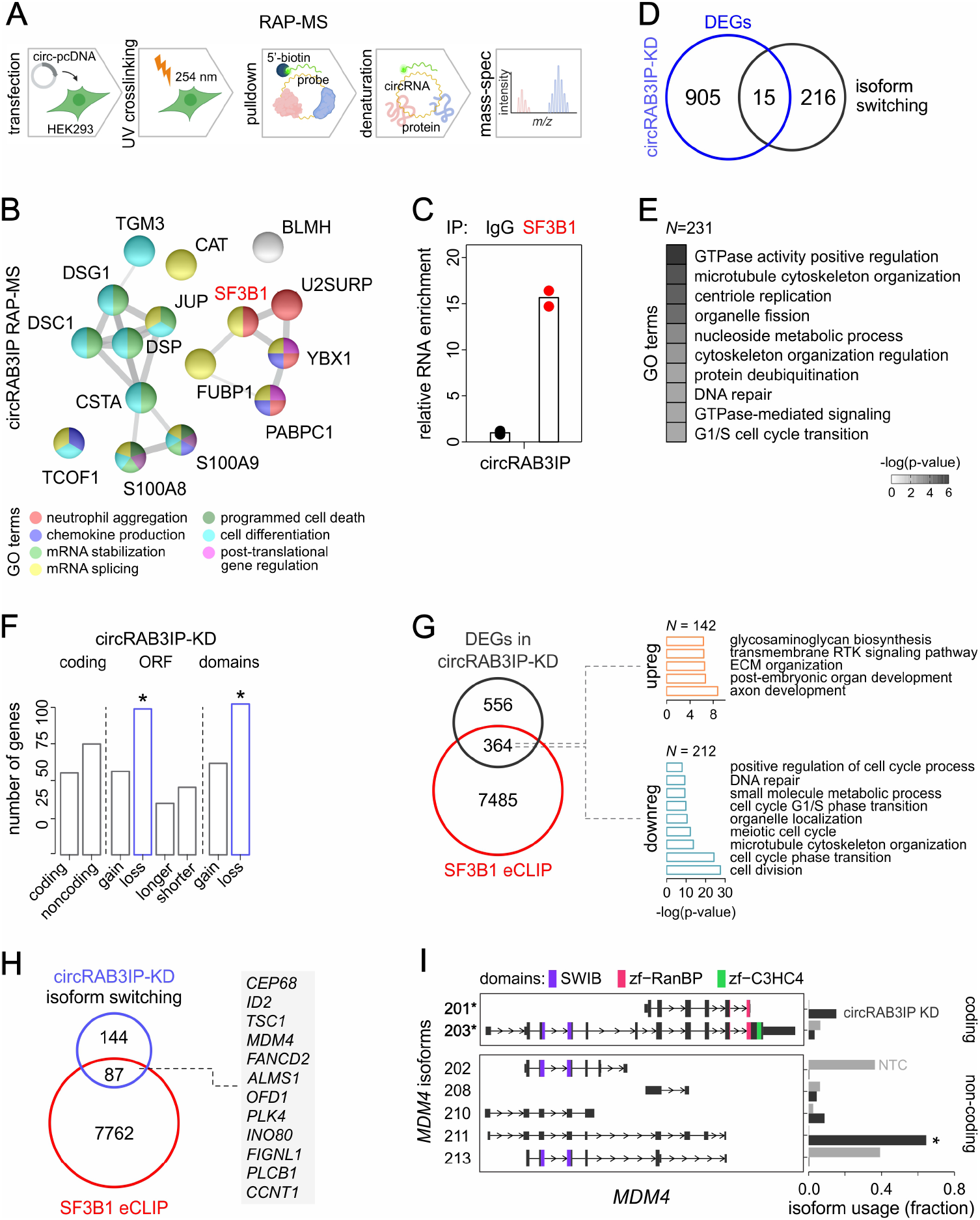
circRAB3IP interacts with SF3B1 and affects RNA splicing. **A,** For RAP-MS, cells are transfected with circRNA-pcDNA3 vectors, UV crosslinked, lysed, and hybridized with biotinylated probes targeting circRNA backsplicing junctions. Following pulldown, circRNA-associated proteins are denatured and analysed by label-free mass spectrometry. **B,** Plot showing circRAB3IP-bound proteins and their interactions inferred from STRING (from three replicates; https://string-db.org/). Color coding reflects GO terms (*key below*) associated with each protein. **C,** Plot showing circRAB3IP enrichment in SF3B1-pulldown experiments in HUVECs normalized to input (±SD, from two replicates). IgG pulldown provides a control. **D,** Venn diagram showing overlap between DEGs and genes displaying isoform usage changes in circRAB3IP-KD data. Overlap was not more than expected by chance; p = 0.6711, hypergeometric test. **E,** GO terms enriched for genes (*N*) with ≥ 30% isoform usage switching in circRAB3IP-KD data. **F,** Bar plot of isoform usage types in circRAB3IP-KD data. *: significant changes with FDR < 0.05. **G,** Venn diagram (*left*) showing the overlap of SF3B1-bound mRNAs with DEGs in circRAB3IP-KD. GO terms (*right*) associated with shared up-/downregulated genes (*orange/blue*). **H,** Venn diagram showing the overlap of SF3B1-bound and differentially spliced mRNAs upon circRAB3IP-KD. *: more than expected by chance; *P* < 0.001, hypergeometric test. Genes linked to cell cycle regulation are listed (*right*; https://gsea-msigdb.org). **I,** Illustration (*left*) of *MDM4* isoforms detected upon circRAB3IP-KD. Functional domains in each isoform (color code, *top*) and coding isoforms (*) are indicated. Bar plot (*right*) showing changes in isoform usage between KD (*black*) and control RNA-seq data (*grey*) for each isoform. *: FDR ≤ 0.001.

circRAB3IP RAP-MS robustly enriched for our circRNA target (**Fig. S3C**) to identify 16 proteins as potential interactors. The majority of these were functionally linked to mRNA splicing and stabilization, chemokine production or cell differentiation (**Fig. 3B**). RAP-MS for circCAMSAP1 returned a catalogue of 27 proteins, most of which could be linked to inflammation, gene expression regulation, and RNA metabolism (**Fig. S3D**). The only interactor common for circRAB3IP and circCAMSAP1 was U2SURP, a known component of the general splicing machinery (Will, 2002). However, the DEGs from our two circRNA-KD experiments showed negligible correlation to DEGs from published U2SURP-KD data from HEK293 cells (R = −0.035 and −0.25 for circRAB3IP- and circCAMSAP1-KD, respectively; De Maio *et al*, 2018). Thus, circCAMSAP1 likely operates via a set of partners that are discrete from those of circRAB3IP.

Of all candidate interactions, we chose to focus on that between SF3B1 and circRAB3IP. This was because SF3B1 is an RBP that has been implicated in mRNA splicing (Sun, 2020), but also in cellular senescence (Yin *et al*, 2019) and aging (Holly *et al*, 2014), and because circRNAs have been shown to affect splicing regulation (Li *et al*, 2015c; Ashwal-Fluss *et al*, 2014). To study the SF3B1-circRAB3IP relationship, we first orthogonally confirmed its association with circRAB3IP in HUVECs via RNA immunoprecipitation (**Fig. 3C**). Next, we reanalyzed our circRAB3IP-KD data to find >200 genes with substantial (>30%) change in isoform usage upon knockdown (**Fig. 3D**). By contrast, circCAMSAP1 depletion induced such changes in just 40 genes. circRAB3IP-related isoform switching is not reflected in differential mRNA expression, as <6% of DEGs also switch isoforms (**Fig. 3D**). Notably, according to GO term analysis, genes displaying differential isoform usage were again linked to cell cycle (G1/S) progression, cytoskeleton reorganization, and centriole regulation (**Fig. 3E**). Almost half of these differentially-spliced genes now expressed isoforms containing fewer functional domains and/or shorter open reading frames (**Fig. 3F**), thus providing an interpretation to how cell cycle deceleration manifests. Those alternative splicing events that are mostly responsible for the observed isoform switching were increased skipping of canonical 5’ and 3’ splicing sites, intron retention, and changes in termination site choice (**Fig. S3E**). Thus, on top of its effect on gene expression levels, an additional regulatory layer involving widespread mRNA processing changes follows circRAB3IP depletion in growing HUVECs.

Finally, we analyzed publicly available SF3B1 eCLIP data (accession no.: ENCSR133QEA) against our circRAB3IP-KD to discover that ~40% of DEGs are also bound by SF3B1 and linked to cell cycle regulation and ECM remodeling (**Fig. 3G**), much like in KD cells (**Fig. 2D,E**). We also analyzed SF3B1 eCLIP against mRNAs that switch isoforms upon circRAB3IP-KD. Again, ~40% of these were also bound SF3B1 (**Fig. 3H**) and contained key cell cycle regulators like *MDM4, ID2, ALMS1, PLK4, FANCD2, CCNT1*, and *PLCB1*. For all of them, strong changes in favor of non-coding/domain-loss isoforms can explain cell cycle effects in cells lacking circRAB3IP (**Figs 3I** and **S3F**). Notably, the negligible overlap between DEGs from circRAB3IP- and published SF3B1-KD experiments (accession no.: GSE88630 and GSE176778; *not shown*) suggests that the circRAB3IP-SF3B1 interaction is selective for the SF3B1 target sub-repertoire linked to cell cycle regulation.

### circRAB3IP induces HUVEC proliferation in a paracrine manner

Given that cell cycle arrest was a prominent feature in both our circRNA-KDs, we reasoned that circRNA overexpression would lead to increased proliferation, in line with what has already been reported for circCAMSAP1 in cancer cells (Zhou *et al*, 2020a; Chen *et al*, 2021; Luo *et al*, 2021; Liu *et al*, 2019). Indeed, MTT assays of the HEK293 cells overexpressing circRAB3IP or circCAMSAP1 that we used in RAP-MS showed significantly accelerated proliferation compared to control cells (**Fig. S4A,B**; circCAMSAP1-, circRAB3IP- and empty vector-overexpressing HEK293 doubling times were 17.4, 22.7 and 28.1 h, respectively).

To study this further, we generated a line allowing for the controllable circRAB3IP production while also negating any negative effects of long-term stable overexpression. We introduced a piggyback construct (pB) into HEK293 cells that efficiently produced circRAB3IP in a doxycycline-induced manner (**Fig. 4A** and **Methods**). Drawing from reports on the packaging and secretion of circRNAs (Li *et al*, 2015b; Dou *et al*,2016), we asked whether this ~2000-fold circRNA excess would end up in extracellular vesicles. Indeed, extracellular vesicles collected from pB-circRAB3IP cultures contained significantly higher circRAB3IP titers than those from unmodified cells (**Fig. 4A**). In fact, blocking vesicle formation leads to circRAB3IP FISH signal accumulation in the cytoplasm of pB-modified cells (**Fig. S4C**).

**Figure 4.**
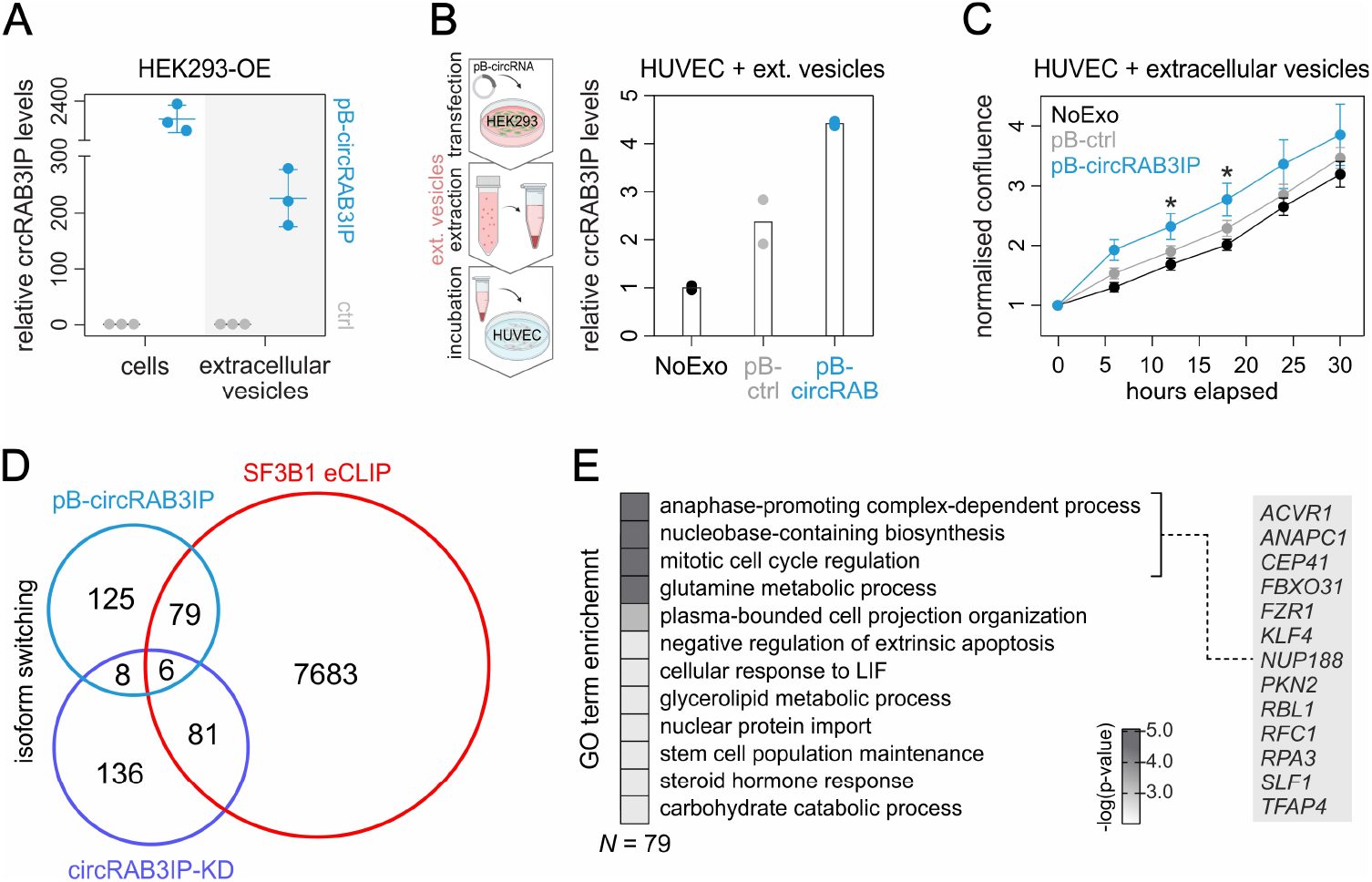
circRAB3IP acts in a cell non-autonomous manner to promote HUVEC proliferation. **A,** Relative circRAB3IP levels (±SD, from three replicates) in empty vector (ctrl) and pB-circRAB3IP cells, and in the respective purified extracellular vesicles. **B,** *Left*: circRNA-pB HEK293 lines were induced for 48h, before extracellular vesicles were purified and added onto growing HUVECs. *Right*: As in panel A, but for HUVECs treated with extracellular vesicles from control (pB-ctrl) and pB-circRAB3IP cells, and expressed as fold-change over non-treated cell levels (NoExo). **C,** Plot showing normalized area confluence (±SD) derived from live-cell imaging of HUVECs stimulated with nontreated, control, and pB-circRAB3IP exosomes from three replicates. *: P_adj_ <0.05, multiple t-testing. **D,** Venn diagram of the overlap between SF3B1 eCLIP targets (*red*) and genes switching isoforms in circRAB3IP-KD (*blue*) or in pB-circRAB3IP exosome-treated cells (*light blue*). **E,** Gene ontology terms associated with the isoform-switching genes in pB-circRAB3IP exosome-treated HUVECs that are also bound by SF3B1. Of these, cell cycle-related genes are listed (*right*).

This setup allowed us to test whether HUVEC proliferation could be affected by higher circRAB3IP titers, given that endothelial cells are not easily amenable for overexpression studies. We collected extracellular vesicles from circRAB3IP-overexpressing or empty vector HEK293 and delivered them onto HUVEC cultures. This resulted in a moderate yet detectable uptake of circRAB3IP (**Fig. 4B**). Next, HUVEC growth rates were monitored over the course of 30 h via automated live-cell imaging. HUVECs showed constantly increasing proliferation rates, despite moderate circRAB3IP uptake (**Fig. 4C**). These experiments highlight the ability of circRAB3IP to modulate cellular growth via paracrine signaling, even when introduced to cells at such low doses.

Since the removal of circRAB3IP leads to apparent reprogramming of gene expression also through splicing changes, we speculated that excess circRAB3IP would induce cell proliferation in a similar manner. We performed RNA-seq in HUVECs treated with circRAB3IP-exosomes to only find negligible changes in gene expression (just 7 genes satisfied a *P*_adj_ < 0.05 threshold; *not shown*). However, following analysis of splicing changes, we discovered >200 transcripts displaying alternative isoform usage (by >20% or more). While negligible overlap to circRAB3IP-KD data was observed, circRAB3IP-treated cells shared 79 transcripts with the list of direct SF3B1 eCLIP targets (**Fig. 4D**). These transcripts were associated with such GO terms as mitotic cell cycle regulation and anaphase-promoting complex-dependent process, and were derived from genes known to exert cell cycle control (**Fig. 4E**). For example, *ANAPC1* and *FZR1* encoding core components of the anaphase-promoting complex (Yamano, 2019), the *KLF4* transcription factor gene whose isoforms have been shown to modulate cell cycle choices (Yang *et al*, 2020) or the known tumor suppressor *RBL1* gene (Liban *et al*, 2017). Changes in isoform usage identified for each of these genes were favorable for cell cycle progression (**Fig. S5**). Therefore, circRAB3IP is required for maintaining homeostatic cell cycle regulation and its depletion or (moderate) overexpression alters this primarily via the isoform modulation of SF3B1-associated transcripts.

## DISCUSSION

Despite a growing body of evidence on the roles of circRNAs in various cancer lines, how they function in primary cells under homeostasis remains largely unexplored. Here, we address this question using human endothelial cells to provide the first functional characterization of circRAB3IP, while also providing a resource for previously-studied circRNAs, like circCAMSAP1. While our motivation for studying this subset of circRNAs originated in their enrichment in nascent RNA, our results showed that circCAMSAP1 and circRAB3IP exhibit functional specificity and distinct interactomes in line with their disparate roles.

circCAMSAP1 has been studied in the context of cancer (Zhou *et al*, 2020a; Chen *et al*, 2021; Luo *et al*, 2021) and chronic inflammation (Liu *et al*, 2019). The former studies found circCAMSAP1 acting as a “sponge” for various microRNAs, whereas the latter implicates its secondary folding in an interaction with PKR. Here, we describe a different role for circCAMSAP1 in cell homeostasis. Its knockdown had overlapping but not fully convergent effects to circRAB3IP depletion, affirming the view of circRNAs as highly specialized molecules carrying out cell type-specific functions. In addition to the senescence-like cell cycle effects in circCAMSAP1-depleted cells, a prominent proinflammatory signature also emerged. This suggests that circCAMSAP1 plays a role in regulating a subset of inflammation-responsive genes, including a considerable fraction of known SASP genes. However, our RAP-MS data did not recover the previously documented interaction of circCAMSAP1 with PKR that is thought to implicate this circRNA in inflammatory regulation (Liu *et al*, 2019). On the other hand, and in line with our analysis of E2F transcription factor motif enrichment in its knockdown-regulated genes, circCAMSAP1 has been shown to influence E2F activity in neurons (Zhou *et al*, 2020a). This could explain how circCAMSAP1 overexpression in HEK293 accelerates also cell cycle progression. However, how its protein interactors, catalogued here by RAP-MS, mediate the functions of circCAMSAP1 remains to be addressed.

circRAB3IP depletion from HUVECs is accompanied by a prominent senescence-like phenotype. circRAB3IP-induced gene expression changes counter homeostatic cell cycle progression and correlate well with the consensus gene expression signature of senescence (Hernandez-Segura *et al*, 2018), as well as with the induction of genes involved in ECM remodeling, another senescence hallmark (Levi *et al*, 2020). Of all the circRAB3IP interactors, the general splicing factor SF3B1 appeared most relevant due to its implication in cell cycle arrest (Dolatshad *et al*, 2015) and senescence (Yin *et al*, 2019). Although the precise nature of the SF3B1-circRAB3IP interplay remains to be determined, its outcome in terms of differential isoform production is central to the effects observed in both the attenuation and acceleration of the cell cycle (e.g., via isoform switching of *RBL1* transcripts). SF3B1 is a core component of the SF3B protein complex that regulates U2-snRNP recruitment to its targets (Zhou *et al*, 2020b) and was recently reported to be guided onto target mRNAs by co-transcriptional association with nucleosomes, rather than by specific nucleotide sequences in the target transcripts themselves (Kfir *et al*, 2015). Given that SF3B1 translocation from the nucleosome onto the nascent transcript is thought to be RNA-dependent, we speculate that circRAB3IP can modulate this association, even at low circRNA titers and for a specific subset of RNA targets. In this scenario, its interaction with U2SURP (which also interacts with circCAMSAP1), a component of U2 snRNP, may be relevant. Such a mode-of-action could explain the well-documented role of splicing in the processes of cellular senescence and tissue aging (Deschênes & Chabot, 2017).

Finally, circRAB3IP can exert its influence on the cell cycle also in a paracrine fashion, despite the very moderate uptake in our system. This opens up the possibility of cell-cell communication via the load carried by the extracellular vesicles exchanged in a given tissue niche. Therefore, our results support the notion that even mild modulation of the levels of a single circRNA can inflict profound changes to gene expression and cellular homeostasis in both a cell autonomous and a cell non-autonomous fashion.

## Author contributions

NJ, KKE, AZ, and AD performed experiments; NJ and CD performed computational analyses; AZ, CD, and AP conceived the study; NJ and AP wrote the manuscript with input from KKE, CD, LK, and TBH.

## Acknowledgements

We wish to thank the Cologne Center for Genomics (CCG) for support with next-generation sequencing, and the Proteomics Core Facility (CECAD, Cologne) for generating all mass spectrometry data. This work was supported by funding from the Deutsche Forschungsgemeinschaft (DFG) via the Priority Program 1935 (Project 313408820) and a Basic Module grant (Project 290613333) awarded to AP, by the Independent Research Fund Denmark and the Novo Nordisk Foundation (Project NNF16OC0019874) awarded to TBH, and by the NRW Stem Cell Network (Grants 3681000801 and 2681101801), the DFG (KU3511/4-1 and /10-1), the Center for Molecular Medicine Cologne (Grant 7102-9530-0005-22) and Koeln Fortune funds awarded to LK.

## Data availability

All RNA-seq data produced in this study are available via the NCBI Gene Expression Omnibus repository.

## METHODS

### Cell culture and circRNA verification

HUVECs from pooled, apparently healthy, donors (passage 4-9; Lonza) were grown in complete Endopan-3 medium kit (PAN-Biotech) at 37°C under 5% CO_2_. HEK293 cells were passaged in DMEM (Sigma) supplemented with 10% FBS (Gibco) and 1% Pen/Strep (Gibco) under the same conditions. For RNAse R treatments, HUVECs were grown in complete media, RNA was extracted via the Direct-Zol RNA purification kit (Zymo) and treated with 1.7 U RNAse R per μg of RNA (Epicenter) for 13 min at 37°C. Fractionation experiments were performed by HUVEC nuclei extraction in a physiological buffer (PB; 100 mM CH_3_COOK, 30 mM KCl, 10 mM Na_2_HPO_4_, 1 mM MgCl_2_, 1 mM Na_2_ATP, 1 mM DTT, 10 mM β-glycerophosphate, 10 mM NaF, 0.2 mM Na_3_VO_4_, 25 U/ml RiboLock RNase Inhibitor, and 1:1000 dilution of cOmplete Protease Inhibitor Cocktail; adjusted to pH 7.4 with KH_2_PO_4_) for 30 min on ice, followed by 5 min centrifugation at 1000 x *g*. Supernatants corresponding to the cytoplasmic fraction were collected in TRIzol, while nuclear pellets washed once more in PB before collection in TRIzol and purification. HUVEC polysome profiling was performed as previously described (Bartsch *et al*, 2018).

### Factory-seq and circRNA detection

Preparations of nascent RNA from ~10^7^ HUVECs were performed as previously described (Melnik *et al*, 2016). Briefly, cells were grown and passaged in complete Endopan-2 medium (Lonza) until 90% confluency and scraped in PB. Nuclei isolation was performed for 30 min in icecold PB supplemented with 0.4% Igepal (Sigma-Aldrich). Lysis efficiency was assessed microscopically. Nuclei were collected by centrifugation and DNase I-treated for 30 min at 37°C (Worthington, 30U per 5 x 10^6^ cells). After DNase I digestion, nuclei were pelleted and lysed in a native lysis buffer (NLB; 40 mM Trisacetate pH 7.4, 2 M 6-aminocaproic acid and 7% sucrose supplemented with 50 U/ml RiboLock RNase Inhibitor and 1:1000 dilution of cOmplete Protease Inhibitor Cocktail) for 20 min on ice. Lysates were further treated with 2U per sample of a Group-III caspase mix (Active human caspases 6, 8, 9, and 10; BioCat) for 30 min under vigorous shaking. Supernatants were collected in TRIzol (Invitrogen) and RNA was isolated using the Direct-Zol RNA purification kit (Zymo). RNA integrity was assessed on a Bioanalyzer and the resulting libraries sequenced to approximately 35 x 10^6^ paired-end reads. Sequencing quality was assessed via FastQC (https://www.bioinformatics.babraham.ac.uk/projects/fastqc/) and reads aligned to the human reference genome (hg19) using STAR (Dobin *et al*, 2013). For detection of circRNAs enriched in nascent RNA, DCC was applied (Cheng *et al*, 2016). Raw read coverage plots were generated using the *Gviz* package (Hahne & Ivanek, 2016).

### circRNA fluorescence *in situ* hybridization

FISH was performed using back-splicing junction-targeting fluorescent probes for circCAMSAP1 and circRAB3IP (listed in **Table S4**) as previously described (Zirkel & Papantonis, 2018). Briefly, HUVECs were grown on coverslips “etched” with 0.1% hydrofluoric acid until 70% confluent and fixed for 15 min (in 10 ml of fixation buffer containing 1 ml 9% NaCl, 2.5 ml 16% paraformaldehyde, 500 μl glacial acetic acid, and 6 ml RNase-free water). Coverslips were then washed in PBS and cells permeabilized in 0.5% Triton-X / 0.5% saponin for 5 min. After washing again in PBS, coverslips were post-fixed in 3.7% formaldehyde for 5 min, washed in PBS and dehydrated in ethanol (70%, 90% and absolute) for 3 min. Hybridization of probes was carried out in hybridization buffer [25% formamide, 2x SSC, 200 ng/μl sheared salmon sperm DNA, 5x Denhardt’s solution, 50 mM phosphate buffer (20 mM KH_2_PO_4_, 30 mM KHPO_4_*2H_2_O, pH 7.0), and 1 mM EDTA] mixed with fluorescent probe (25 ng/μl) in a 9:1 ratio. The hybridization mix was denatured at 90°C for 10 min, quenched on ice for 2 min and added to the coverslips. Hybridization was performed overnight in the dark at 37°C in a humid chamber. The next day, coverslips were washed three times in 2x SCC, once in RNase-free water, and mounted on glass slides with ProLong Gold Antifade pre-supplemented with DAPI (Invitrogen). Imaging was performed on a Leica DMi8 platform using a 63x magnification objective (oil), and images exported via the LASX software (Leica).

### siRNA-mediated circRNA knockdown and analysis

Approximately 35 x 10^4^ HUVEC cells were seeded per each 60 mm plate 1 day before transfection. Each siRNA (180 pmol) was mixed with Opti-MEM (Gibco) and Lipofectamine RNAiMAX according to manufacturer’s instructions (Thermo Fisher). Cells were washed once in PBS and incubated in 2.5 ml of Opti-MEM and 500 μl of transfection mix for 4 h, after which the medium was replaced with complete Endopan medium. 48 h post-transfection, cells were collected in TRIzol (Invitrogen) and RNA extraction was performed using the Direct-Zol RNA purification kit (Zymo). RNA integrity was verified with Bioanalyzer, and samples were sequenced to at least 25 x 10^6^ paired end reads. Sequencing quality check was performed using FastQC and reads were aligned to human reference genome (hg19) with default STAR aligner settings (Dobin *et al*, 2013). Aligned reads quantification was performed with *featureCounts* (Liao *et al*, 2014), selecting for uniquely aligned and properly matched read pairs (selected options --primary -p -B -C). Between-sample normalization of raw quantified reads was performed using the RUVs option of RUVseq (Risso *et al*, 2014) that estimates factors of unwanted variation using as reference those genes in the replicates for which the covariates of interest remain constant. Differentially expressed genes (DEGs) were estimated via DESeq2 (Love *et al*, 2014), wherein genes with FDR <0.01 and absolute (log_2_) fold-change ≥0.6 were considered significantly differentially expressed. The resulting DEGs (deposited in GEO alongside raw data) were used as an *a priori* defined list for gene set enrichment analysis (GSEA; Subramanian *et al*, 2005) to assess the overrepresentation of GO terms in the RNA-seq datasets and produce network plots via Cytoscape (Franz *et al*, 2016). Read counts of mapped RNA-seq were normalized per million mapped reads (RPM) and plotted using *ngs.plot* (Shen *et al*, 2014) to visualize signal enrichment along genes of pathways chosen from the GSEA Molecular Signatures Database (MSigDB IDs: TNFα via NFkB signaling, M5890; IFNγ response, M5913; E2F targets, M5925; G2/M checkpoint, M5901). GO term analysis for gene sets without assigned expression data (eCLIP and isoform switching genes) was performed using Metascape (Zhou *et al*, 2019a). Isoform switch analysis was performed via *IsoformSwitchAnalyzeR* (v1.12; (Vitting-Seerup & Sandelin, 2019), whereby raw RNA-seq data is first pseudoaligned to hg19 using *Salmon* (v1.2; Patro *et al*, 2017) and the resulting count matrices were used in to find genes displaying no significant change in expression levels, but rather in transcript usage. After default filtering steps and statistical testing for significance, genes which display at least 20% (for circRNAs delivered by extracellular vesicles) or 30% change in isoform usage (for circRNA-KD) were considered for downstream analyses.

### circRNA overexpression constructs

Expression vectors for circCAMSAP1 and circRAB3IP were generated by amplifying genomic DNA containing the two circRNA-generating exons, the intron in between them and ~500 and ~100 bp of the up- and downstream flanking introns. PCR amplified fragments were inserted into pcDNA3 or piggyBAC (pB) vectors via appropriate restriction digests and ligation. To facilitate circRNA production, artificial inverted repeats were generated around the circRNA exons by PCR amplification of ~360 bp of the upstream flanking intron and its insertion in an inverted orientation downstream of the circRNA exons through a restriction digest. For Northern blots and RNase R treatment of cells transfected with pcDNA3 overexpression vectors, 5 μg RNA were digested with 1 U RNase R (Epicentre) per μg RNA in a total reaction volume of 10 μl for 10 min at 37°C. After this, samples were prepared for agarose blotting by addition of 20 μl loading buffer to 5 μg RNase R-treated or untreated purified RNA. Samples were then loaded onto a 1% agarose gel containing 3% formaldehyde and 1x MOPS and run at 75 V in 1x MOPS for ~3 hs after which RNA was transferred onto a Hybond N+ membrane (GE Healthcare) overnight. RNA was then UV crosslinked to the membrane and prehybridized in Church buffer (0.158 M NaH_2_PO_4_, 0.342 M Na_2_HPO_4_, 7% SDS, 1 mM EDTA, 0.5% BSA at pH 7.5) for 1 h. The membrane was probed with a 5’ radioactively labelled DNA oligonucleotide (60-mer probe) at 55 °C overnight and washed twice in 2x SSC, 0.1% SDS for 10 min at 45°C before exposure on a Phosphoimager screen for data collection. Cell proliferation of pB transfected HEK293 cells was monitored via MTT assays. In brief, triplicates of ~2,000 cells were seeded in one well of a 96-well plate. On the next day, the medium was replaced with 100 μl fresh medium plus 10 μl of a 12 mM MTT stock solution (Invitrogen), and incubated at 37°C for 3 h. Subsequently, all but 25 μl of the medium was removed from the wells, and formazan dissolved in 50 μl DMSO was mixed thoroughly with the cells and incubated at 37°C for 10 min. Samples were then mixed again and absorbance was read at 530 nm. Measurements were taken at 24, 48, 72 and 96 h post-seeding and background levels subtracted.

### RNA antisense-purification coupled to mass spectrometry (RAP-MS)

Approximately 4 x 10^8^ HEK293 were transfected with either circCAMSAP-pcDNA3 or circRAB3IP-pcDNA3 vectors via CaPO_3_ transfection. Briefly, cells were transfected 48 h prior to the experiment with a mix containing the plasmid, 2.5M CaCl_2_, and HEBS (50 mM HEPES, 280 mM NaCl, 1.5 mM Na_2_HPO_4_ at pH 7.05 adjusted using NaOH). An additional dish of HEK293 was transfected with control GFP-pcDNA3 to monitor transfection efficiency. RAP-MS was performed essentially as previously described (McHugh & Guttman, 2018) with 55-mer sense and antisense biotinylated probes designed to target circCAMSAP1 or circRAB3IP. Cells were UV-crosslinked at 254 nm (Stratalinker 1800; 0.8 J/cm^2^), scraped in ice-cold PBS, collected by centrifugation, and counted to ensure 2 x 10^8^ cells per capture reaction. Cell lysis was performed for 10 min on ice in total cell lysis buffer (10 mM Tris-HCl pH 7.5, 500 mM LiCl, 0.5% dodecyl-maltoside, 0.2% SDS, 0.1% sodium deoxycholate, 1x cOmplete Protease Inhibitor Cocktail, and 1000 U of RiboLock RNAse Inhibitor). During cell lysis, samples were passed through 26G needles 5 times and then sonicated (Bioruptor microtip sonicator, 30% amplitude) for 15 cycles with 5 sec “on” / 10 sec “off” settings. Following sonication, lysates were treated with 30 U DNase I (Worthington) for 20 min at 37°C and precleared for 30 min at 52°C with 1.2 ml of streptavidin Roti MagBeads beads (Carl Roth) per capture reaction. Beads which were previously washed in 10 mM Tris-HCl pH 7.5 and hybridization buffer (10 mM Tris-HCl pH 7.5, 5 mM EDTA, 500 mM LiCl, 0.5% DDM, 0.2% SDS, 0.1% sodium deoxycholate, 4 M urea, 2.5 mM TCEP). After preclearing, 2 nmol of each probe was denatured at 85°C, quenched on ice, added to the lysates, and incubated at 52°C for 4 h with intermittent mixing (12 sec “on” / 9 sec “off”, 900 rpm). For probe capture, a fresh aliquot of streptavidin beads was prepared as above and added to the samples for another 30 min at 52°C. Beads were then washed 6x in hybridization buffer for 5 min at 52°C with intermittent mixing. After the final wash, beads were resuspended in benzonase elution buffer (20 mM Tris-HCl pH 8.0, 0.05% NLS, 2 mM MgCl_2_, and 0.5 mM TCEP) and hybridized probes eluted from the beads using 120 U Benzonase (Sigma) for 2 h at 37°C with intermittent mixing. Supernatants were collected and treated with 5 mM DTT for 30 min at 55°C, and subsequently with 40 mM CAA for 30 min at room temperature, centrifuged for 10 min at 20,000 x *g*, transferred to a new tube, and stored at −20°C until preparation for mass spectrometry. The final sample preparation and mass spectrometry analysis was carried out at the Proteomics Core Facility in Cologne, Germany. To assess the efficiency of each RNA pull-down, input and eluted RNA samples were collected during the experiment (as described by McHugh & Guttman, 2018), resuspended in NLS elution buffer (20 mM Tris-HCl pH 8.0, 10 mM EDTA, 2% NLS, 2.5 mM TCEP) and treated with 1 mg/μl of Proteinase K for 1 h at 55°C. RNA was then isolated via phenol:chloroform extraction, precipitated with ethanol, and reverse transcription was carried out using the SuperScript™ II Reverse Transcriptase according to manufacturer’s instructions (Invitrogen). qPCR reactions were run using the qPCRBIO SyGreen Mix Separate-ROX (NIPPON).

### RNA immunoprecipitation coupled to qPCR (RIP-qPCR)

HEK293 cells grown to confluence in complete media were harvested by gentle scaping in PBS, counted to ensure ~2 x 10^7^ cells per IP, pelleted, and lysed in ice-cold polysome lysis buffer (100 mM KCl, 5 mM MgCl_2_, 10 mM HEPES-NaOH pH 7.0, 1 mM DTT, 200 U/ml RNaseIn, 1x PIC, 0.5% NP-40) for 30 min. Cell lysis was facilitated by passing the lysates through 26G needle 10 times during the incubation on ice, and by 6 cycles of sonication (30 sec “on”/30 sec “off”, low input) on a Bioruptor Pico (Diagenode). Lysates were then treated with 40 U DNase I (Worthington) per IP for 30 min and pelleted by centrifugation to remove incompletely lysed cells. Input (5%) samples were taken from supernatants and stored in TRIzol (Invitrogen) until RNA extraction. The rest of the supernatants was subjected to overnight immunoprecipitation with 10 μg of IgG (Milipore, 12-371B; 1 μg/μl) or SF3B1 antibody (Santa Cruz, sc-514655; 0.2 μg/μl). Next day, 30 μl of protein-G Dynabeads (Invitrogen) were prewashed with 1 ml of NT-2 buffer (250 mM Tris-HCl pH 7.4, 750 mM NaCl, 5 mM MgCl_2_, 0.25% NP-40) and incubated for 1 h with 10 μg/IP of mouse bridging antibody (Active Motif; 1 μg/μl) at 4°C with end-to-end rotation. Beads were next washed 3 times in NT-2 buffer, added to the lysates, and incubated for 4h at 4°C with end-to-end rotation. Lysates were next washed 6 times in 1 ml of NT-2 buffer for 3 min/wash on end-to-end rotor to ensure removal of unbound material. After the last wash, beads were resuspended in TRIzol (Invitrogen) and RNA extracted using the Direct-zol RNA kit (Zymo). Reverse transcription was carried out using the SuperScript™ II Reverse Transcriptase (Invitrogen) and qPCRs using the qPCRBIO SyGreen Mix Separate-ROX (NIPPON).

### Extracellular vesicle purification and transfer experiments

HUVECs and pB-circRNA transfected HEK293 were grown in media supplemented with exosome-depleted (Life Technologies). HEK293 were induced with 3 μg/ml doxycycline (Sigma) and 400 μg/ml of G418 (Sigma). After 48 h, cells were collected in TRIzol for qPCR analysis of overexpression, and media collected on ice to purify extracellular vesicles through a series of centrifugation steps (Livshts *et al*, 2015). Briefly, first round of centrifugation was at 750 x *g* for 10 min to remove leftover cells from the medium. Supernatants were then collected and spun at 1,500 x *g* for 10 min, followed by 14,000 x *g* for 35 min and 100,000 x *g* for 2 h to separate microvesicles and exosomes. Pellets containing extracellular vesicles were then washed in PBS (Sigma) and subjected to another centrifugation at 100,000 x *g* for 1h. Pellets carrying extracellular vesicles were resuspended in 200 μl PBS and stored short-term at −80°C. For assessment of circRNA enrichment in HUVECs, exosomes were used for RNA extraction (Total Exosome RNA & Protein Isolation Kit, Invitrogen), according to the manufacturer’s instructions and qPCR analysis as above. For extracellular vesicle transfer onto HUVECs, cells were grown in complete media supplemented with purified extracellular vesicles for 24 h. HUVECs were then seeded and monitored for growth in an IncuCyte S3 platform (Sartorius) for 30 h. For RNA-seq analysis of extracellular vesicle-treated HUVECs, cells were washed 2x in PBS and collected in TRIzol to extract RNA.

### Statistical tests

Assessment of correlation between RNA-seq datasets was performed using the Pearson’s correlation core function in R (https://www.r-project.org/); Student’s t-tests, Fisher’s exact test, and nonlinear regression fits were performed using Prism (https://www.graphpad.com/scientific-software/prism/). Unless stated otherwise, any data with a *P*-value <0.05 were deemed as significant.

