## Supplemental Figs S1-S5 and Tables S1-S4 for "circRAB3IP modulates cell proliferation by reorganizing gene expression and mRNA processing in a paracrine manner"

- This supplement contains **Figures S1-S5** and **Tables S1-S4**.

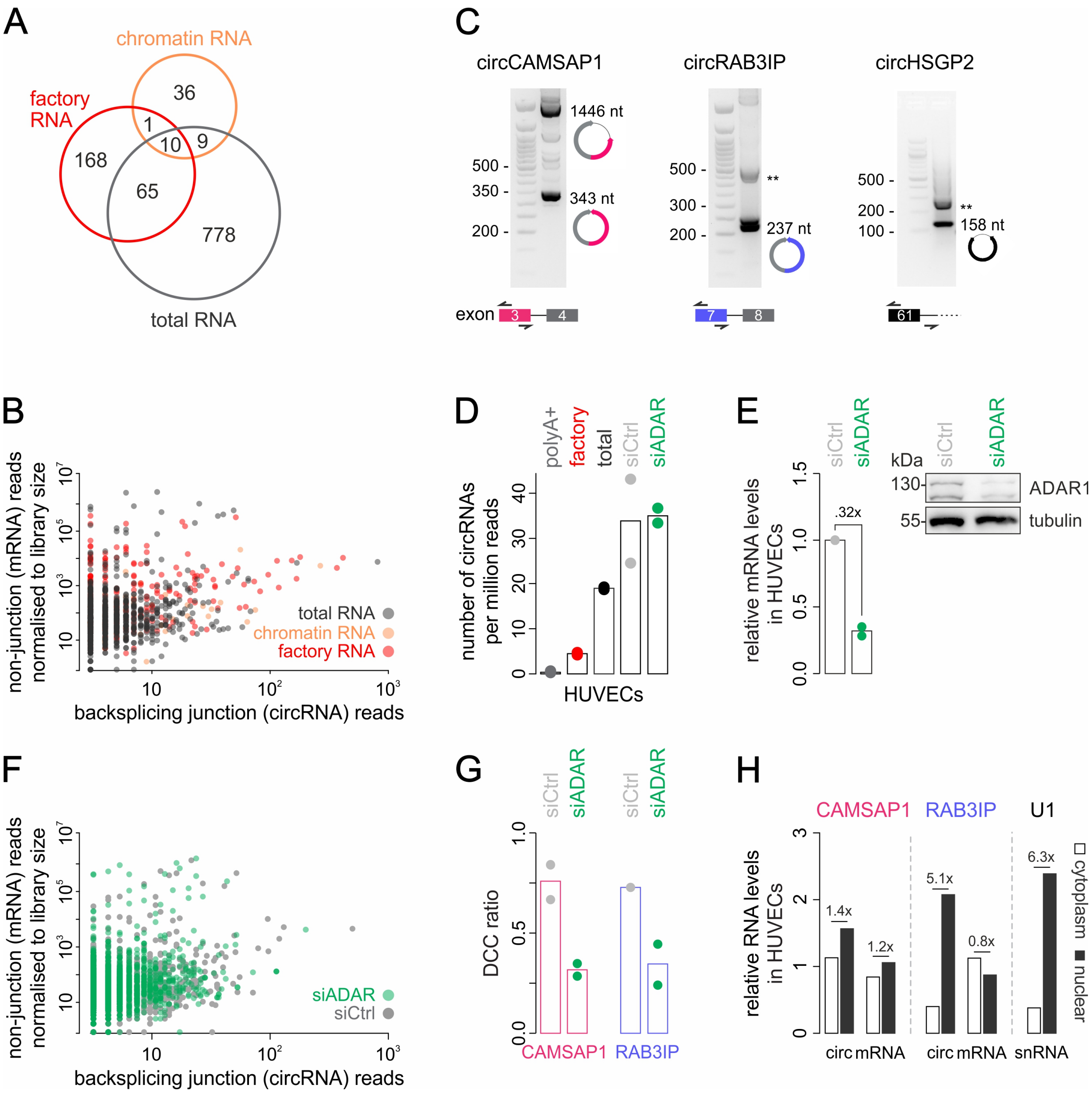


**Figure S1. Verification of circRNA identities and ADAR1 sensitivity.**

**A,** Venn diagram showing detection of circRNAs in factory (*red*), chromatin-associated (*orange*) or total RNA fraction from HUVECs (*dark grey*).

**B,** Scatter plot of the number of reads mapping to circRNA backsplicing junctions plotted against reads from the same exons mapping to non-junction reads in the three RNA fractions from panel A.

**C,** Electrophoresis profiles showing circCAMSAP1 (*left*), circRAB3IP (*middle*), and circHSPG2 amplification (*right*) by PCR using divergent primers (*bottom*) with sizes indicated (*cartoons*). Size ladders are run in the first lane of each gel. **: PCR artefacts.

**D,** Bar plot showing the number of circRNAs detected in polyA-selected (*grey*), nascent "factory" (*red*), total RNA fractions (*black*), and in control (*light grey*) and ADAR-KD libraries (*green*) from HUVECs.

**E,** Bar plots (*left*) showing fold-reduction in *ADAR1* levels upon knockdown from two replicates in HUVECs. Western blotting (*right*) confirms ADAR1 knockdown and β-tubulin levels provide a control.

**F,** As in panel B, but for control (*light grey*) and ADAR1-KD libraries (*green*) from HUVECs.

**G,** Bar plots showing changes in DCC ratios for circCAMSAP1 and circRAB3IP upon ADAR-KD (two replicates). **H,** Bar plot showing relative enrichment of circCAMSAP1, circRBA3IP or their mRNA counterparts in subcellular fractions as determined by RT-qPCR (from two replicates). U1 snRNA levels provide a control.

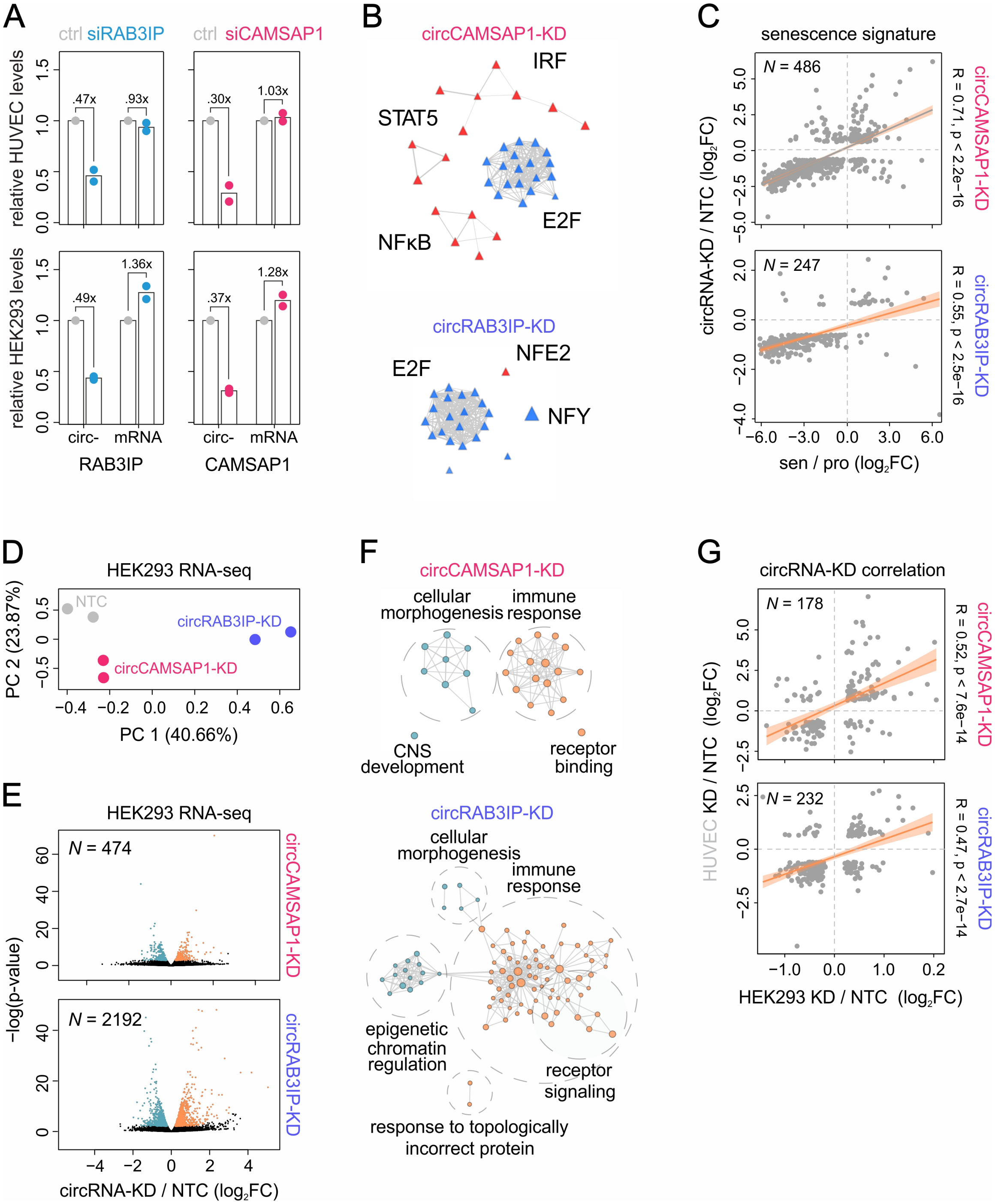
A
ctrl siRAB3IP
ctrl siCAMSAP1
relative HUVEC levels
relative HEK293 levels
.47x
.93x
.30x
1.03x
.49x
1.36x
.37x
1.28x
circ- mRNA
RAB3IP
circ- mRNA
CAMSAP1
B
circCAMSAP1-KD
IRF
STAT5
E2F
NFκB
circRAB3IP-KD
E2F
NFE2
NFY
C
senescence signature
N = 486
N = 247
circRNA-KD / NTC (log2FC)
sen / pro (log2FC)
R = 0.71, p < 2.2e−16
circCAMSAP1-KD
R = 0.55, p < 2.5e−16
circRAB3IP-KD
D
HEK293 RNA-seq
NTC
circRAB3IP-KD
circCAMSAP1-KD
PC 2 (23.87%)
PC 1 (40.66%)
E
HEK293 RNA-seq
N = 474
N = 2192
−log(p-value)
circRNA-KD / NTC (log2FC)
circCAMSAP1-KD
circRAB3IP-KD
F
circCAMSAP1-KD
cellular morphogenesis
immune response
CNS development
receptor binding
circRAB3IP-KD
cellular morphogenesis
immune response
epigenetic chromatin regulation
receptor signaling
response to topologically incorrect protein
G
circRNA-KD correlation
N = 178
N = 232
HUVEC KD / NTC (log2FC)
HEK293 KD / NTC (log2FC)
R = 0.52, p < 7.6e−14
circCAMSAP1-KD
R = 0.47, p < 2.7e−14
circRAB3IP-KD

**Figure S2. circCAMSAP1- and circRAB3IP-KD trigger transcriptional changes in HEK293 cells.**

**A,** Bar plots showing the fold-reduction in circRAB3IP and circCAMSAP1 levels upon knockdown in HUVECs (*top*) or HEK293 (*down*; from two replicates). Changes in mRNA levels provide a control.

**B,** Network plot of regulatory transcription factor enrichment for the genes differentially expressed upon circCAMSAP- (*top*) or circRAB3IP-KD (*bottom*) inferred via overrepresentation analysis.

**C,** Scatter plot correlating circCAMSAP1- (*top*) and circRAB3IP-KD RNA-seq data (*bottom*) with the gene expression signature for IMR90 replicative senescence. Pearson correlation coefficients (R) and their associated p-values for overlapping DEGs (*N*) are indicated.

**D,** PCA plot of RNA-seq replicates of circCAMSAP1- and circRAB3IP-KD in HEK293 cells.

**E,** Volcano plots showing fold changes ($\log_2$) for up-/downregulated genes (*orange*/*blue*) following HEK293 circCAMSAP1- (*top*) or circRAB3IP-KD (*bottom*) at a significance cut-off set of $-\log_{10}(p\text{-}value) \geq 1.3$.

**F,** Gene set enrichment analysis for genes differentially expressed in circCAMSAP1- (*top*) and circRAB3IP-KD (*bottom*) in HEK293 (absolute $\log_2$ fold change $\geq 0.6$ and $p_{adj} \leq 0.05$). Clusters of related terms were merged and labelled by the group-dominant gene ontology terms.

**G,** As in panel C, but correlating HEK293 with HUVEC circRNA-KD DEGs.

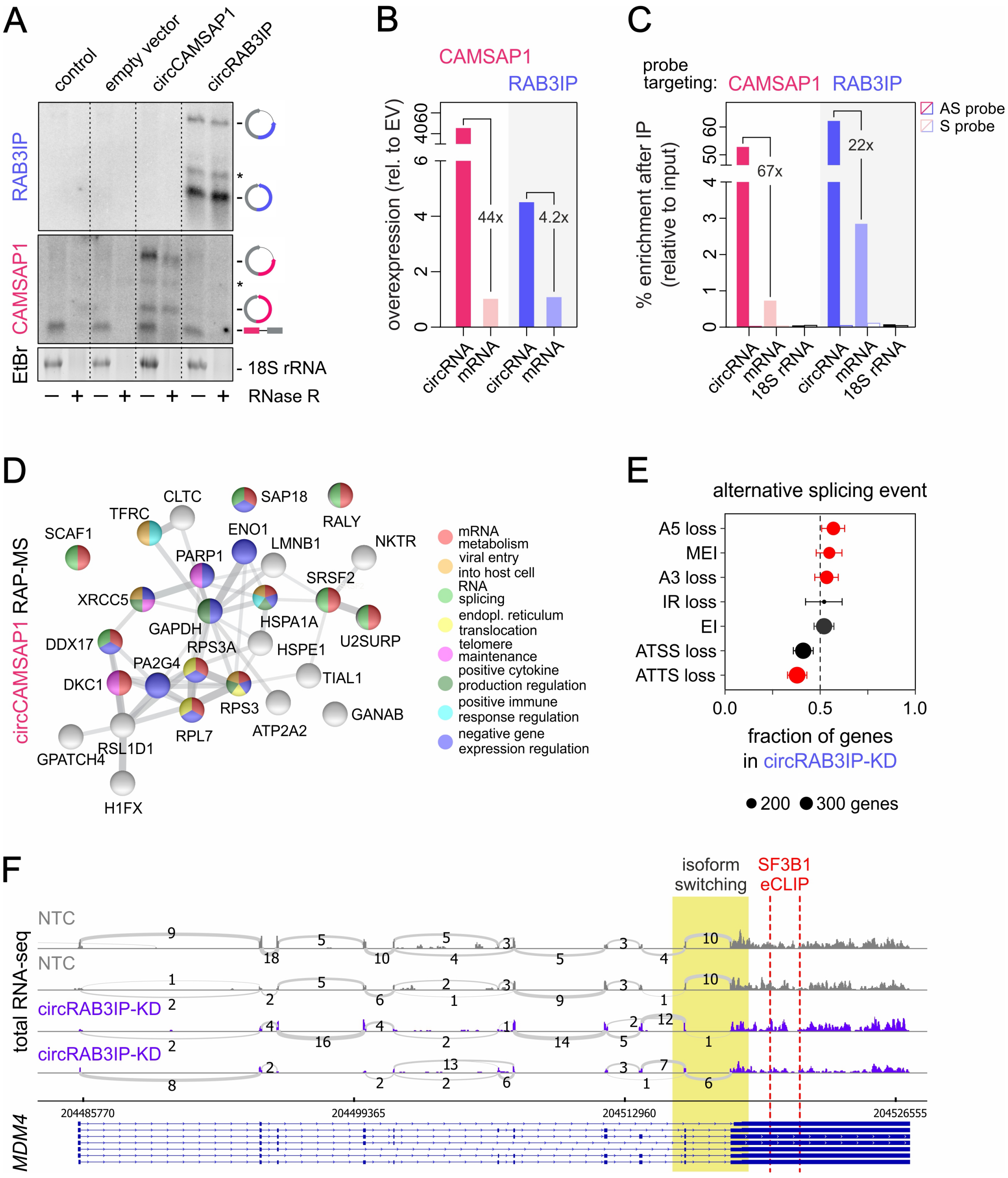
A
control
empty vector
circCAMSAP1
circRAB3IP
RAB3IP
CAMSAP1
EtBr
18S rRNA
RNase R
B
CAMSAP1
RAB3IP
overexpression (rel. to EV)
44x
4.2x
circRNA
mRNA
C
probe targeting:
AS probe
S probe
% enrichment after IP (relative to input)
67x
22x
18S rRNA
D
circCAMSAP1 RAP-MS
mRNA metabolism
viral entry into host cell
RNA splicing
endopl. reticulum translocation
telomere maintenance
positive cytokine production regulation
positive immune response regulation
negative gene expression regulation
E
alternative splicing event
A5 loss
MEI
A3 loss
IR loss
EI
ATSS loss
ATTS loss
fraction of genes in circRAB3IP-KD
200
300 genes
F
isoform switching
SF3B1 eCLIP
total RNA-seq
NTC
circRAB3IP-KD
204485770
204499365
204512960
204526555
MDM4

**Figure S3. circRNA RAP-MS and circRAB3IP-KD effects on isoform usage.**

**A,** Northern blot for untransfected (control), empty vector (EV), and pcDNA3-circRNA transfected HEK293 with or without RNase R treatment. 18S rRNA levels provide a control. *: artefactual circRNA concatamers.

**B,** Bar plots showing circCAMSAP1 and circRAB3IP overexpression levels in HEK293 (from two replicates) normalized relative to *YWHAZ* and represented as fold-change over empty vector (EV) cells. mRNA levels of the respective host genes provide a baseline.

**C,** Bar plots showing per cent circCAMSAP1 and circRAB3IP enrichment after RAP-MS in HEK293 (from two replicates) normalized relative to input. Fold-enrichments over *CAMSAP1* or *RAB3IP* mRNA are shown. 18S enrichment as well as enrichments following the use of a sense probe (S) provide a negative control.

**D,** STRING interaction network of the proteins co-purifying with circCAMSAP1 in RAP-MS experiments. Proteins enriching for a particular Gene Ontology term (*key on the right*) are color-coded.

**E,** Genome-wide changes of alternative splicing events (with ≥ 10% change) in circRAB3IP- KD data. Dot size indicates the number of genes and significant changes (*red*) are supported by an FDR < 0.05. A5, alternative 5' donor site; A3, alternative 3' donor site; MEI, multiple exon inclusion; ES, exon inclusion; IR, intron retention; ATSS and ATTS, alternative transcription start and transcription termination sites, respectively.

**F,** Sashimi plot for the *MDM4* gene in control (NTC) and circRAB3IP-KD HUVECs. The number of normalized read counts for each exon-exon junction are indicated. The exons involved in isoform switching (*yellow rectangle*) and positions bound by SF3B1 (*red dashed line*) are highlighted.

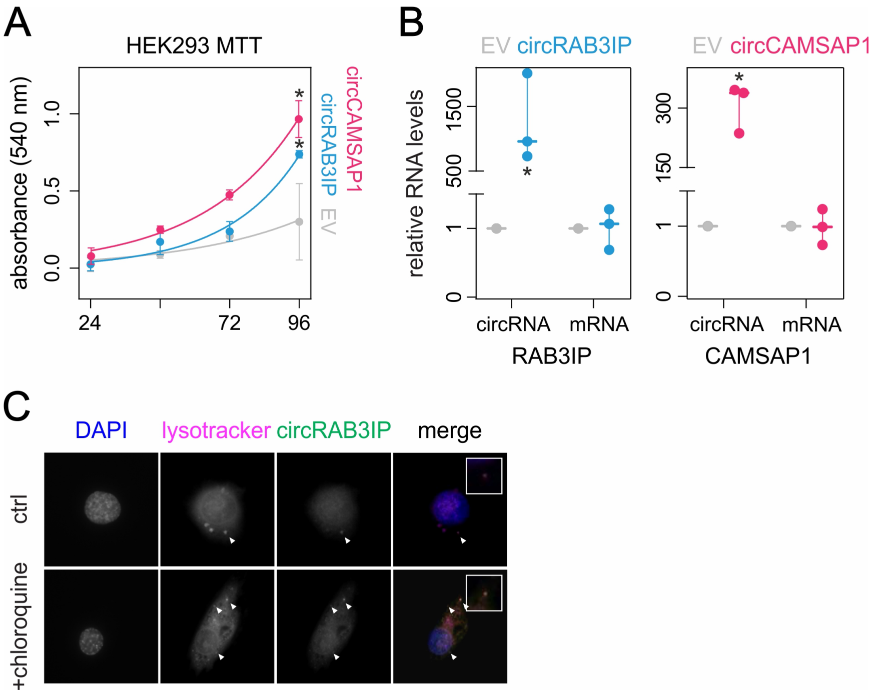


**Figure S4. Effects of circRNA overexpression in human cells.**

**A,** MTT assays (±SD from three replicates) under stable circRAB3IP (*blue*), circCAMSAP1 (*magenta*) or empty vector overexpression (*grey*). Relative absorbance was plotted relative to the 24-h time point, and doubling times were 17.4 h for circCAMSAP1-, 22.7 h for circRAB3IP- and 28.1 h for EV-overexpression. *: significantly different to EV control; $p < 0.05$, unpaired two-tailed Student's t-test.

**B,** Plot showing circRAB3IP (*left*) and circCAMSAP1 overexpression levels (*right*) in the HEK293 cells used in panel A (three replicates) normalized to *YWHAZ* and represented as fold-change over empty vector (EV) levels. *: significantly different to EV control; $p < 0.05$, unpaired two-tailed Student's t-test.

**C,** Representative circRAB3IP RNA FISH images of HUVECs treated or not (ctrl) with 10 µM chloroquine and counterstained with DAPI. circRAB3IP FISH signal accumulation is indicated (*arrowheads*).

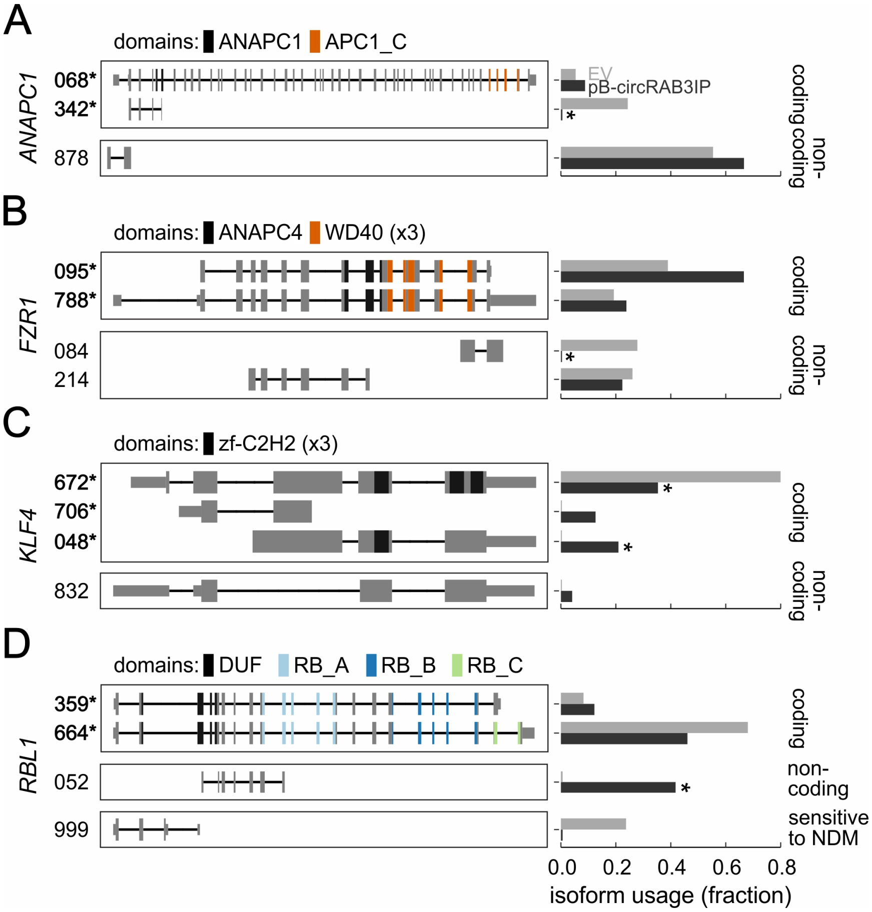


**Figure S5. Examples of isoform switching following paracrine pB-circRAB3IP treatment of HUVECs.**

**A,** Illustration (*left*) showing detected *ANAPC1* isoforms. Functional domains in each isoform (color code, *top*) and coding isoforms (*) are indicated. Bar plot (*right*) showing usage changes between pB-circRAB3IP-treated (*black*) and control HUVEC RNA-seq data (*grey*) for each isoform. *significantly different to control: FDR ≤ 0.001.

**B,** As in panel A, but for *FZR1* isoforms.

**C,** As in panel A, but for *KLF4* isoforms.

**D,** As in panel A, but for *RBL1* isoforms.

**Table S1.** List of circRNA junctions, their respective read counts and DCC ratios from factory, chromatin and total RNA-seq experiments, as well as from ADAR-KD in HUVECs (data provided as an *.xlsx* file).

**Table S2.** List of peptides identified via RAP-MS experiments in HEK293 cells overexpressing circCAMSAP1 or circRAB3IP. Usual MS contaminants are marked in grey (data provided as an *.xlsx* file).

**Table S3.** Oligonucleotides used as PCR primers, siRNAs or cloning primers

| PCR targets | forward primer (5'-3') | reverse primer (5'-3') |
|---|---|---|
| circRAB3IP | GTTCTGGAGGCTGTGGAAAA | GACACGTCCTGTCCATTGTG |
| *RAB3IP* mRNA | GAGGCTTTCAAAAGTGCGAG | TGTTTTTCTGCTGTTGCCTG |
| circCAMSAP1 | CCCTGATGATGGCCTACACT | CTCAGGGATGTTATCTTGTTGAT |
| *CAMSAP1* mRNA | ATTGCCCAGAGCAGATGAAA | ATACAGACTGTCGGCCATCG |
| *NFKBIA* mRNA | AGACCTGGCCTTCCTCAACTTCC | AGTCTCGGAGCTCAGGATCACAG |
| *IL32* mRNA | CGAATGCACCAGGCCATAGA | GACATCACCTGTCCACGTCC |
| *CCL20* mRNA | TATTGTGGGCTTCACACGGC | GATTTGCGCACACAGACAAC |
| *TNFAIP3* mRNA | TCGACAGAAACATCCAGGCCAC | TGTCCTGAACGCCCCACATGT |
| *U1* snRNA | CAGGGGAGATACCATGATCACGAAG | GGTCAGCACATCCGGAGTGCAATGG |
| **cloning targets** | **Sequence (5'-3')** | |
| circCAMSAP1-F-pcDNA (*Bam*HI) | GATCGGATCCCATGGGTGTTCACCCTCTGT | |
| circCAMSAP1-F-pB (*Mlu*I) | GATCACGCGTCATGGGTGTTCACCCTCTGT | |
| circCAMSAP1-R-pcDNA (*Not*I) | GATCGCGGCCGCTATTCACTGGCCAGGTGTGG | |
| circCAMSAP1-R-pB (*Pac*I) | GATCTTAATTAAGACTGGTCCCAAATGAAGGG | |
| circCAMSAP1-InvF-pcDNA (*Apa*I) | GATCGGGCCCATGGGTGTTCACCCTCTGT | |
| circcircCAMSAP1-InvF-pB (*Pac*I) | GATCTTAATTAAGAGCAACTAACAACGTGAAAAGG | |
| circCAMSAP1-InvR-pcDNA (*Not*I) | GATCCCGCGGCCGCGAGCAACTAACAACGTGAAAAGG | |
| circCAMSAP1-InvR-pB (*Spe*I) | GATCACTAGTCATGGGTGTTCACCCTCTGT | |
| circRAB3IP-F-pcDNA (*Not*I) | ATCGGCGGCCGCACAGTTCAGCCAGTCTCAGT | |
| circRAB3IP-F-pB (*Mlu*I) | ATCGACGCGTACAGTTCAGCCAGTCTCAGT | |
| circRAB3IP-R-pcDNA (*Xho*I) | ATCGCTCGAGCTGCAACCATAACATTTCAA | |
| circRab3IP-R-pB (*Pac*I) | ATCGTTAATTAACTGCAACCATAACATTTCAA | |
| circRAB3IP-InvF-pcDNA (*Apa*I) | TCGAGGGCCCACAGTTCAGCCAGTCTCAGT | |
| circRAB3IP-InvF-pB (*Pac*I) | ATCGTTAATTAACGCTTTACAAAACCAAAACT | |
| circRAB3IP-InvR-pcDNA (*Xho*I) | TCGACTCGAGAAATTGAACCACTCTTGGTG | |
| circRAB3IP-InvR-pB (*Spe*I) | ATCGACTAGTACAGTTCAGCCAGTCTCAGT | |
| **siRNA targets** | **Sequence (5'-3')** | |
| circCAMSAP1 | GATCAACAAGATAACATCCCT | |
| circRAB3IP | GGAGGACCAAAGCTGACTTAT | |
| ADAR1-1 | CGCAGAGUUCCUCACCUGUTT | |
| ADAR1-2 | GGAUGCAAAUCAAGAGAAATT | |

**Table S4.** Oligonucleotides used as probes in circRNA FISH, RAP-MS, and Northern blots.

| circRNA FISH | sequence (5’-3’) |
|---|---|
| circCAMSAP1 ex.3-2 | TCTCTGAGGTCCTCAGGGATGTTATCTTGTTGATCCAGAACACCATGGCATCCTC |
| circRAB3IP ex.8-7 | GAATTCATTATACAAGGATAAGTCAGCTTTGGTCCTCCGCATTCAACTGCAGAAG |
| circHSGP2 ex.61-intron | CGCTGCTGTGGCTCCACTCTGTACCTGGGACGCTGGGGTCACCAGCTGGCTCAGG |
| **RAP-MS** | **sequence (5’-3’)** |
| circCAMSAP1-Sense | TCTCTGAGGTCCTCAGGGATGTTATCTTGTTGATCCAGAACACCATGGCATCCTC |
| circCAMSAP1-AS | GAGGATGCCATGGTGTTCTGGATCAACAAGATAACATCCCTGAGGACCTCAGAGA |
| circRAB3IP-Sense | GAATTCATTATACAAGGATAAGTCAGCTTTGGTCCTCCGCATTCAACTGCAGAAG |
| circRAB3IP-AS | CTTCTGCAGTTGAATGCGGAGGACCAAAGCTGACTTATCCTTGTATAATGAATT |
| **Northern blot** | **sequence (5’-3’)** |
| circRAB3IP | AAAGGACACGTCCTGTCCATTGTGGGCTCATCCTTCCACAATCGGAATTCATTATAC AAG |
| circCAMSAP1 | TGCGCGACTGAGGTCGGACTCTGTCACGGGGGTGTCATCACTCTCCATC |
